# Fetal manipulation of maternal metabolism is a critical function of *Igf2* imprinting

**DOI:** 10.1101/2023.04.19.537510

**Authors:** Jorge Lopez-Tello, Hannah E. J. Yong, Ionel Sandovici, Efthimia Christoforou, Esteban Salazar-Petres, Rebecca Boyland, Tina Napso, Miguel Constancia, Amanda N. Sferruzzi-Perri

**Affiliations:** Centre for Trophoblast Research, Department of Physiology, Development and Neuroscience, University of Cambridge, Cambridge CB2 3EG, United Kingdom; Singapore Institute for Clinical Sciences, Agency for Science, Technology and Research, Singapore; Department of Obstetrics and Gynaecology and National Institute for Health Research Cambridge Biomedical Research Centre, Cambridge CB2 0SW, United Kingdom; Wellcome-MRC Institute of Metabolic Science and Medical Research Council Metabolic Diseases Unit, University of Cambridge, Cambridge CB2 0QQ, UK; Royal Devon and Exeter Hospital NHS Trust, Barrack Rd, Exeter EX2 5DW, United Kingdom

## Abstract

Maternal-offspring interactions in mammals are mainly characterised by cooperation, but also conflict. Over evolutionary time, the fetus has evolved to manipulate the mother’s physiology to increase nutrient transfer through the placenta, but these mechanisms are poorly characterized. The imprinted *Igf2* (insulin-like growth factor 2) gene is highly expressed in mouse placental cells with endocrine functions. Here, we show that in the mouse, deletion of *Igf2* in these cells leads to impaired placental endocrine signalling to the mother, but remarkably does not result in changes in placental morphology, growth or size. Mechanistically, we find that *Igf2* via defective production of hormones, including prolactins, is essential for the establishment of the insulin-resistance state during pregnancy, and the appropriate partitioning of nutrients to the developing fetus. Consequently, fetuses are growth restricted and hypoglycemic, due to impaired placental glucose transfer from the mother to the fetus. Furthermore, *Igf2* loss from placental endocrine cells has long-lasting effects on offspring adiposity and glucose homeostasis in adult life. Our study provides long-sought compelling experimental evidence for an intrinsic fetal manipulation system, which operates in the placenta to modify maternal metabolism and resource allocation to the fetus, with consequences for offspring metabolic health in later life.

## Main

Maternal-offspring interactions are largely viewed as cooperative, as both the mother and the fetus have an interest in the other’s wellbeing and survival. Pregnancy enhances metabolic demands on the mother to increase nutrient availability for fetal growth and development (Napso et al., 2018). However, the fetus will seek to extract more resources than the mother is prepared to give, as parents and offspring do not share all genes and future maternal reproduction can be negatively impacted if too much investment is placed in the current offspring (Haig, 1993). In the context of pregnancy, this genetically-driven ‘conflict’ over maternal resources has its origins with the first placental mammals around 150 million years ago (Killian et al., 2001; Wood and Oakey, 2006). Genomic imprinting, which co-evolved with placentation, and placental hormone production represent two important mechanisms thought to mediate fetal ‘manipulation’ of maternal physiology for the benefit of the fetus (Fowden and Moore, 2012).

Most studies to date have shown key roles for imprinted genes in regulating the placental supply of nutrients to the fetus, or in the regulation of fetal resource acquisition through actions in the mother, where these genes are also expressed (Constância et al., 2004; Cassidy and Charalambous, 2018). However, direct evidence for a role of placental imprinting or hormones as fetal manipulation systems of maternal resources is currently lacking.

### Fetal hypoglycemia and impaired maternal insulin-resistance response

*Igf2* is a paternally-expressed gene, with the maternal gene copy silenced by epigenetic mechanisms (DeChiara et al., 1991). We employed a mouse model, based on the Cre-loxP technology in which dams carried offspring with either a deletion of the paternally-inherited *Igf2* allele in the placental endocrine cells (referred to as UE litters, with diminished *Igf2* production from the endocrine cells in whole litters), or deletion of the maternally-inherited and silent *Igf2* allele (referred to as control litters, with normal *Igf2* levels in whole litters) (Fig 1A). In litters of mixed genotypes, we have previously shown that this approach leads to an absence of *Igf2* specifically in the endocrine cells of the placenta, indicating that *Igf2* is imprinted in placental endocrine cells (Aykroyd et al., 2020). In litters where all the placentas are UE, IGF2 expression is, as expected, mostly absent in the placental junctional zone by immuno-fluorescence (Fig 1A-C) and reduced at the mRNA level in dissected samples with a degree of labyrinth zone contamination (Fig S1A). Using this strategy, we principally aimed at measuring the impact of loss of *Igf2* from the placental endocrine cells on maternal metabolism, and its consequences for fetal development (Fig 1A-C).

**Figure 1.**
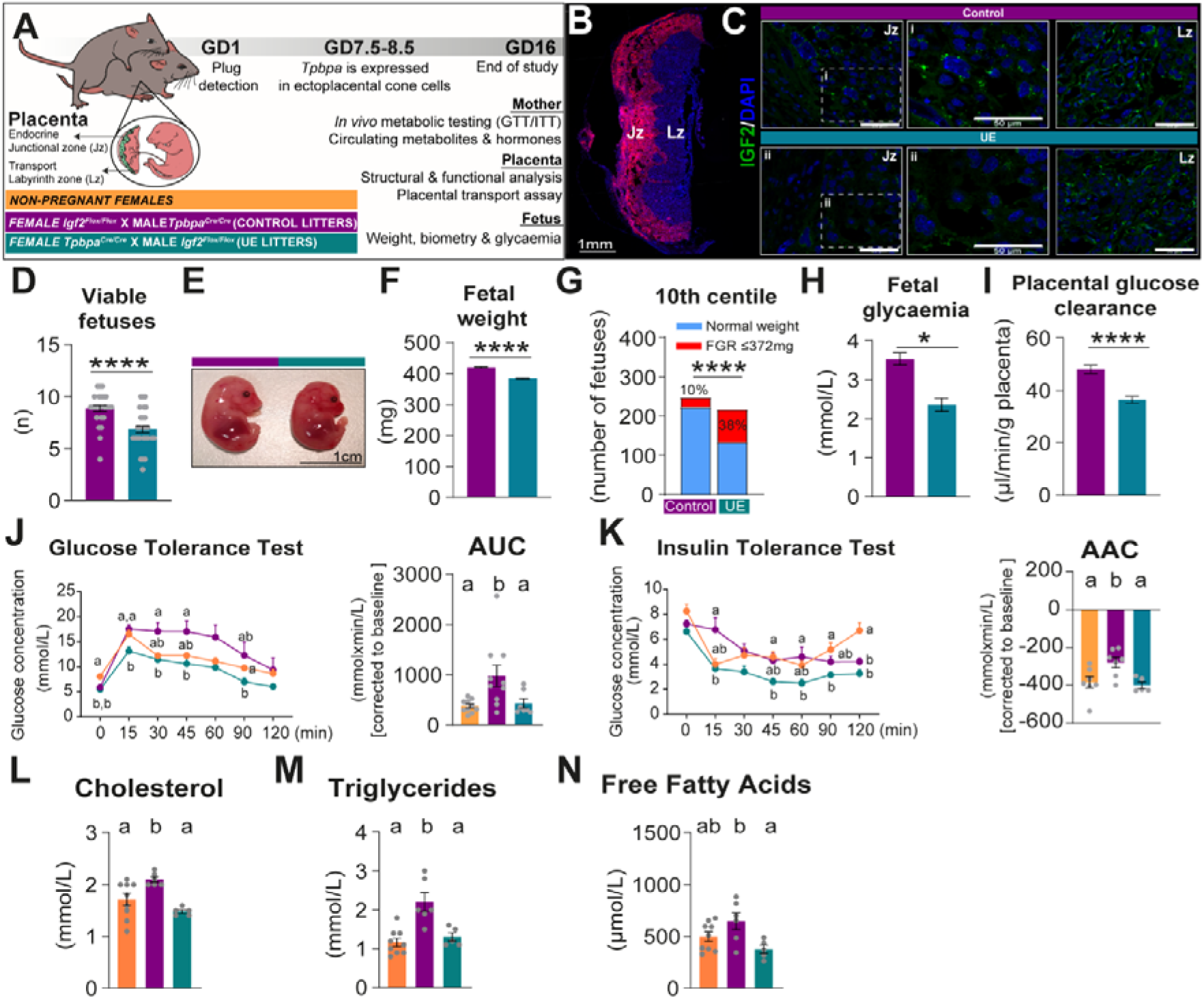
**Reduced *Igf2* in the placental endocrine cells is associated with reduced litter size, fetal growth, failed maternal acquisition of glucose intolerance and insulin resistance during pregnancy, and decreased circulating maternal lipids at GD16**. (A) Schematic of experimental design and mating strategy to disrupt *Igf2* in the placental endocrine zone. (B) TdTomato-Cre fluorescent reporter showing that Cre is exclusively active in the placental endocrine zone (Junctional zone, Jz) and not in the transport region (Labyrinth zone, Lz). (C) Representative IGF2 protein staining in the Jz and Lz for each experimental group. Scale bars are 50 µm. (D) Number of viable fetuses per litter by genotype (n=28-32 litters/group). (E) Representative image of fetuses from each experimental group and absolute fetal weight(n=28-32 litters/group). (F-G) Proportion of fetuses considered as having fetal growth restriction (defined as fetal weight below the 10^th^ centile cut-off – 372mg of the control group, n=28-32 litters/group). (H) Fetal blood glucose concentration (assessed in two randomly selected fetuses/litter, n=8-9 litters/group). (I) *In vivo* placental transport of ^3^H-methyl-D glucose (MeG) relative to placental weight (n=7-8 litters/ group). (J-K) Intraperitoneal glucose and insulin tolerance tests (GTT and ITT). Inserts show area under glucose curve (AUC) and area above insulin curve (AAC). (n=5-10 dams/group/test). (L-N) Concentrations of cholesterol, triglycerides and free (non-esterified) fatty acids in maternal circulation (n=5-9 dams/group/test). Data presented as mean±SEM with or without individual data points shown. Data analysed with Mann Whitney test or one-way ANOVA with Tukey post-hoc tests for mean comparisons, as appropriate. GTT and ITT data were analysed by two-way ANOVA with repeated measures. Feto-placental data were analysed by repeated measures ANOVA (MIXED model), considering the mother as the subject and the genotype the main factor, with litter size added as a covariate. *p<0.05; ***p<0.001; ****p<0.0001. Different letters (a,b,c) indicate significant differences between groups by ANOVA. Scale bars are indicated inside each image.

Exclusive loss of *Igf2* in the endocrine cells of the placental junctional zone led to reduced litter size and symmetrical fetal growth restriction, with a 9% reduction in the average fetus weight at gestational day (GD)16, and increased number of fetuses below the 10^th^ centiles (Fig 1D-G and Fig S1B-G). Alterations in fetal growth were not coupled with a change in placental growth, size, or cellular composition, but resulted, in a decrease in placental efficiency in UE conceptuses (Fig S2A-E). In the absence of placental morphological features that could explain the fetal growth restriction and since glucose is the principal substrate driving fetal growth and energy metabolism, we then measured maternal-fetal parameters of glucose homeostasis. Placenta glucose clearance was reduced by 24% *in vivo* and circulating glucose was significantly lower in UE fetuses compared to control by 33% (Fig 1H-I). Once fetal weight is accounted for, no change in fetal glucose accumulation was observed (Fig S1H-I). Expression of glucose transporters (*Slc2a1* and *Slc2a3*) in the dissected placental labyrinth transport zone was unchanged (Fig S2F).

A major mechanism by which adequate levels of glucose are delivered to the fetus includes the transient state of insulin resistance of maternal tissues that occurs naturally during pregnancy (Musial et al., 2016). *In vivo* metabolic tolerance testing revealed that dams carrying UE litters failed to become glucose intolerant and insulin resistant (Fig 1J-K). Glucose tolerance and insulin sensitivity thresholds were instead at similar levels to non-pregnant controls. Consistent with their retention of insulin sensitivity, dams with UE litters also did not increase their circulating levels of cholesterol, triglycerides and free fatty acids (non-esterified fatty acids, NEFAs) that were observed in pregnant controls; thus, maintaining similar levels to those observed in non-pregnant females (Fig 1L-N).

### Placental and maternal contributions to metabolic dysfunction during pregnancy

To inform on the metabolic dysfunction observed in UE dams and investigate if, and how, loss of *Igf2* affects placental endocrine function, we first prepared primary cultures of placental endocrine cells from control and UE litters at GD16 and analysed the conditioned media using qualitative LC-MS/MS (Fig 2A). A total of 1,408 proteins in the conditioned media were detected (Fig 2B), including known secreted proteins - placental lactogen/prolactin family members (e.g. PRL3B1, PRL8A8, PRL7A1) and pregnancy-specific glycoproteins (e.g. PSG16, PSG21, PSG23) and IGFBPs (e.g. IGFBP6, IGFBP7) (Napso et al., 2021). Uniprot gene ontology analysis focused on the 290 proteins that were detected in one group, but not the other (i.e., 125 unique proteins in UE, and 165 unique proteins in controls), identified biological terms related to metabolic processes that comprised 52% and 68% of the unique proteins in the control and UE secretomes, respectively (Fig 2C and Table S1). Uniprot analysis also revealed additional terms, including hormone/estrogen biosynthetic and reproductive processes (Table S1). Of note, several proteins have been implicated in modulating insulin resistance/sensitivity [e.g. TNFA and semaphorins (Akash et al., 2018; Lu and Zhu, 2020), absent in UE but present in controls], with other proteins showing a qualitative gain or loss in UE, which include key regulators of lipid homeostasis [e.g. FGFR1, APOA4 (Wu et al., 2015; Geng et al., 2020)], glucose metabolism [e.g. LEPR, FUCA1 (Borges et al., 2019; Lu et al., 2016)] and β-cell proliferation [e.g. PRL2C1, FGFR1(Baeyens et al., 2016; Kolodziejski et al., 2020)].

**Figure 2.**
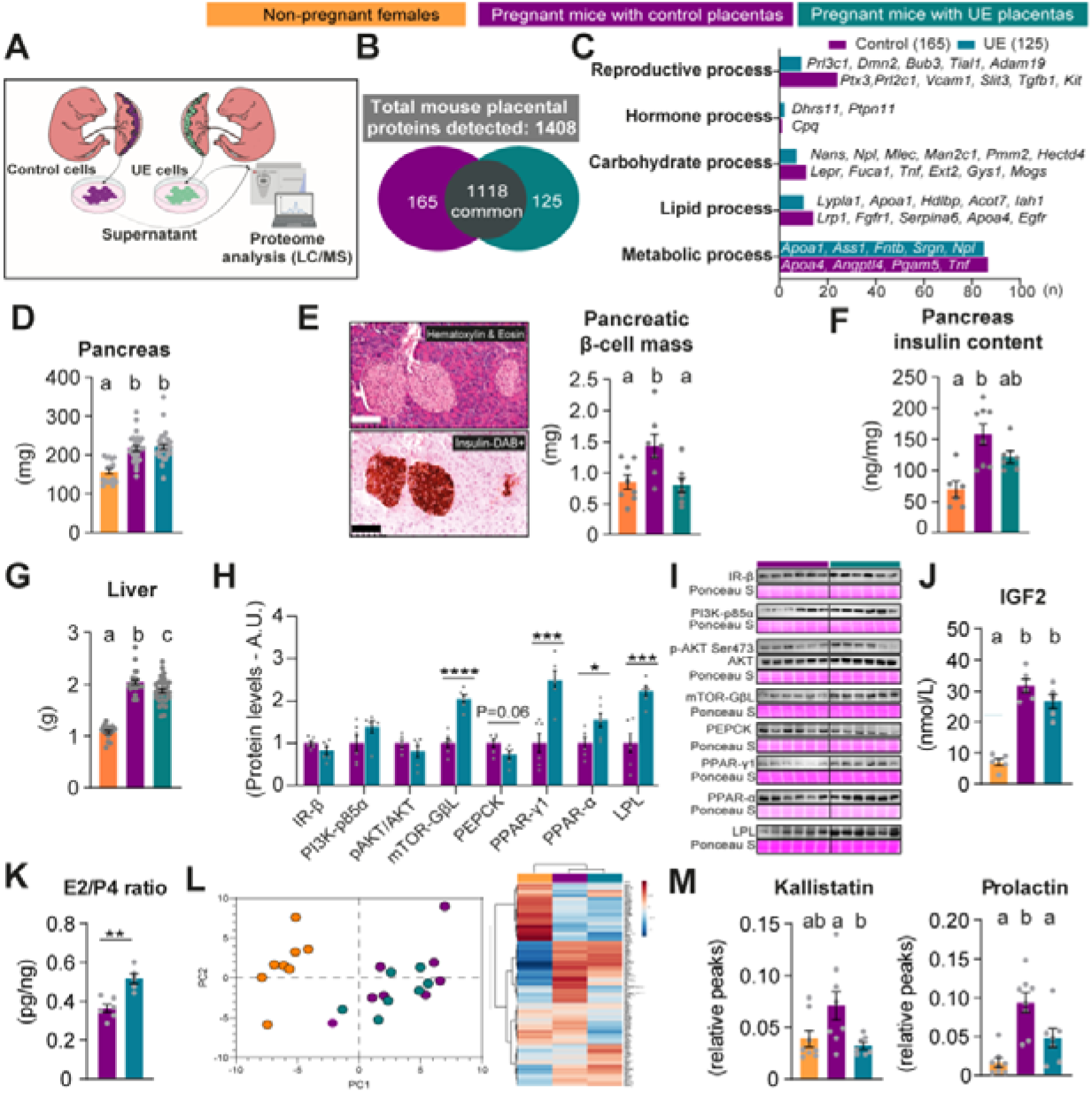
Loss of *Igf2* in placental endocrine cells alters placental secretome and disrupts maternal pancreatic and hepatic adaptation during pregnancy and maternal circulating hormones on GD16. (A) Schematic of experimental design for placental proteomics. Liquid chromatography mass spectrometry (LC-MS) was performed on supernatants derived from conditioned medium of cultured primary cells from dissected junctional zones of control and UE placentas. Need to note that separated primary cultures are not pure (hence, there is potential contamination of labyrinth zone, which may limit detection of junctional zone proteins). (B) Number of placental proteins detectable by LC-MS/MS from control and UE placental secretomes (n=3 litters/group). (C) Relevant pathways identified by Uniprot gene ontology analysis of proteins uniquely present in the control and UE secretomes. On the x axis, n=number of proteins for each gene ontology term. To facilitate the visualisation of the graph, gene names were used instead of protein names. (D) Absolute pancreatic mass (n=16-29/group). (E) Representative histological staining of pancreatic β-cell islets (insulin-containing cells are DAB positive) and quantitated pancreatic β-cell mass (n=7-8/group). Scale bar is 100μm. (F) Pancreas insulin content measured by ELISA (n=6-8/group). (G) Absolute liver mass (n=14-30 per group). (H) Relative levels of proteins involved in glucose and lipid metabolism in maternal liver (n=6/group). (I) Immunoblots of the proteins shown in H with respective Ponceau staining as loading control. (J) Circulating concentrations of IGF2 in maternal plasma (n=5/group). (K) Ratio of estradiol (E2) to progesterone (P4) in maternal plasma detected by ELISA (n=5-6/group). (L) Principal component (PC) analysis plot and heat map of proteins detected in maternal plasma by LC-MS/MS (n=7-8/group). (M) Relative levels of kallistatin and prolactin detected in maternal plasma by LC-MS/MS (n=7-8/group). Data presented as mean±SEM with individual data points shown. Student unpaired t-tests or Mann Whitney test for two groups analysis or ANOVA with Tukey post-hoc tests for mean comparisons were performed as appropriate. *p<0.05; **p<0.01; ***p<0.001; ****p<0.0001. Different letters (a,b,c) indicate significant differences between groups by ANOVA.

Maternal insulin resistance is normally compensated by an increase in pancreatic insulin secretion, which is achieved at least, in part, by an expansion of the β-cell mass during pregnancy (Salazar-Petres and Sferruzzi-Perri). While maternal pancreas weight increased as it should in pregnancy (Fig 2D), β-cell mass expansion failed to occur in UE dams (Fig 2E). Pancreatic insulin content also increased to a lesser extent in dams with UE litters (UE values are intermediate between non-pregnant and pregnant dams with control litters; Fig 2F).

The liver, an insulin-responsive organ important for whole body glucose and lipid handling, did not fully increase in weight in dams with UE litters during pregnancy (values were intermediate between non-pregnant and pregnant control females, Fig 2G). Lipid handling proteins - LPL, an enzyme that mediates the clearance of lipids by hydrolysing them so they can be rapidly taken up by tissues (Essaji et al., 2013), and PPARs (γ,α), important for fatty acid metabolism (Montaigne et al., 2021), and glucose homeostasis (Berthier et al., 2021) - were all increased in the liver of UE dams (Fig 2H-I). mTORC1 protein GβL, which is activated by insulin (Kim et al., 2003; Ong et al., 2016), was also significantly elevated in the liver of UE dams, and there was a tendency for PEPCK, which is inhibited by insulin and is rate-limiting for gluconeogenesis, to be lower in the liver of UE dams (Fig 2H-I). Although there was an absence of deregulation of canonical markers of hepatic insulin signalling (i.e. IR/AKT/PI3K; Fig 2H-I), these findings are consistent with the inability to develop insulin resistance and increase circulating lipids in pregnancy, and are suggestive of a decrease in hepatic glucose production.

To further elucidate the mechanisms that lead to failure in acquiring the pregnancy-associated insulin resistance and β-cell mass expansion in UE dams, we performed unbiased quantitative LC-MS/MS on plasma and investigated the levels of selected maternal circulating hormones. We found no difference in IGF2 concentrations, which are extremely low in adult mice, between control and UE pregnant dams (Fig 2J). Although IGF2 is increased ∼3-fold in pregnancy compared to non-pregnant state, these findings rule out a direct contribution of circulating IGF2 to the metabolic phenotype observed in UE dams. Circulating estradiol to progesterone ratio was greater in dams carrying UE litters, compared to dams carrying control litters (Fig 2K and Fig S3A-B), suggesting perturbations in steroid hormonal production. Proteomic analysis identified 180 proteins in the maternal plasma and these demonstrated a clear segregation between the samples from pregnant and non-pregnant dams, with more subtle differences in clustering between pregnant dams carrying UE compared to control litters (Fig 2L). Of the 180 proteins, 80 were also part of the 1,408 proteins detected in the placental secretome (regardless of placenta genotype), suggesting a placental contribution to the levels measured in the circulation (e.g. SERPINA6, CP, ITIH4, LIFR). Two proteins in the circulation were differentially abundant between the two pregnant groups, namely prolactin, a hormone with roles in metabolic regulation, including insulin sensitivity and β-cell proliferation (Baeyens et al., 2016), and kallistatin (SERPINA4), a serine proteinase inhibitor that is elevated in the circulation of diabetics and correlates with lipid levels (Jenkins et al., 2010) - both proteins being significantly decreased in UE relative to control dams (Fig 2M). The relative abundance of 6 additional proteins in the circulation of UE dams were intermediate between non-pregnant and pregnant control dams, including CFB (Fig S3C-H), which is involved in glucose-insulin handling (Coan et al., 2017). Other proteins like VTNC, which is correlated with hepatic insulin sensitivity (Cao et al., 2018), were significantly elevated in the circulation of UE dams compared to non-pregnant (pregnant controls were intermediate between non-pregnant and UE dams; Fig S3I-K). Several of the proteins that displayed a different pattern of expression in the pregnant UE relative to the non-pregnant and pregnant control dams were among those detected in the placental secretome (e.g. PRL, SERPINA, VTN, UBE2N, GPX1, and APO; Fig 2B), suggesting that the placenta contributes to their differential levels in the circulation (see Table S2 for full list of proteins detected in mouse plasma).

Together, these results show that the likely mediators of maternal body composition changes and other metabolic adaptations related to insulin resistance, lipid and endocrine profile and β-cell mass expansion in UE dams, could comprise a small set of proteins secreted from the UE placenta into the maternal circulation.

### *In utero* placental programming of metabolic dysfunction observed in adult offspring

Direct evidence that perturbation of placental imprinting can programme offspring for metabolic disease in later life is lacking. Taking advantage of our unique placental-specific genetic manipulation, we evaluated the metabolic health and body composition of UE and control litters that were fed, from weaning, either a normal chow diet or a high sugar-high fat diet (HSHFD) (Fig 3A). There was no difference in the weight of the offspring during lactation and following weaning between control and UE litters, with food intake after weaning comparable to their diet-matched control counterparts (Fig S4).

**Figure 3.**
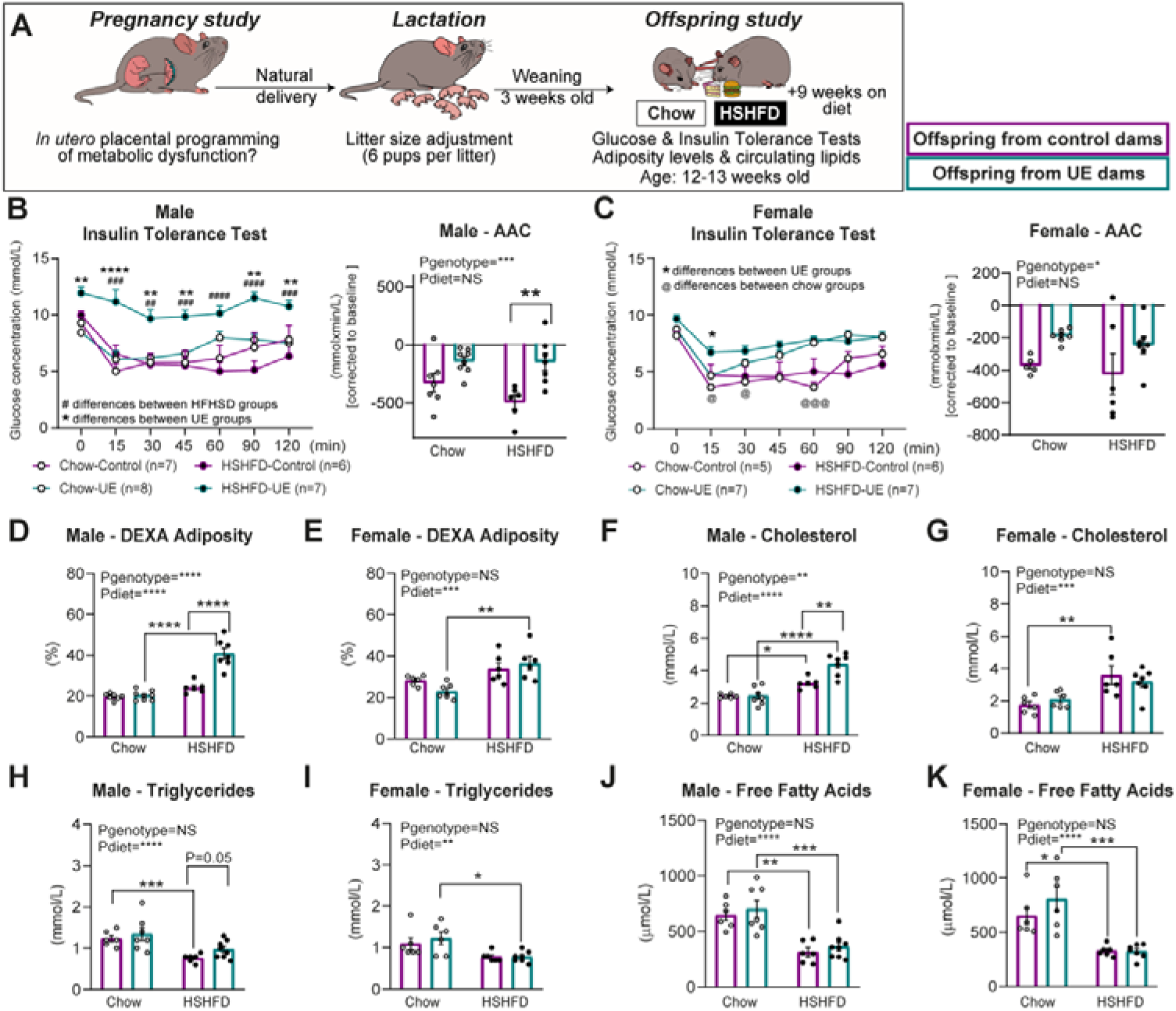
Offspring with placental junctional zone *Igf2* loss have impaired insulin tolerance independent of diet and increased adiposity and altered circulating lipids on a high sugar high fat diet (HSHFD) (A) Schematic of experimental design for offspring follow-up. (B-C) Intraperitoneal insulin tolerance tests were performed on male and female offspring on chow and HFHS diets at 12 weeks of age. Inserts show area above insulin curve (AAC) (n=5-8/group/sex). (D-E) Adiposity as measured by DEXA scans in male and female offspring on chow and HSHFD at 13 weeks of age (n=6-8/group/sex). (F-K) Circulating concentrations of cholesterol, triglycerides and free fatty acids in male and female offspring on chow and HFHSD at 13 weeks of age (n=6-8/group/sex). Data presented as mean±SEM with individual data points shown. Two-way ANOVA was performed with Tukey post-hoc test for mean comparisons. *p<0.05; **p<0.01; ***p<0.001; ****p<0.0001. Different symbols (*, @, #) indicate significant differences between groups by ANOVA.

Notably, UE offspring were insulin resistant, regardless of the postnatal diet or sex, when compared to controls (Fig 3B-C). Glucose tolerance was however, unaffected in UE offspring, fed with chow or HSHFD (Fig S5). While there was no difference in adiposity between UE and control offspring on a chow diet, both UE male and female offspring showed a greater gain of adiposity on a HSHFD (∼20% for UE for both sexes and ∼5% and 10% for male and female controls; Fig 3D-E). Furthermore, UE male offspring were ∼2-times fatter (% adiposity) than control males when they were fed a HSHFD.

Circulating lipids (cholesterol, triglycerides and NEFAs) levels were not different between UE and control offspring on a chow diet, regardless of sex and genotype (Fig. 3F-K). HSHFD significantly increased circulating cholesterol in the offspring. However, this was exacerbated in UE males who had elevated concentrations compared to HSHFD control males (Fig 3F-G). The HSHFD also reduced circulating triglycerides, but this was less pronounced in UE male offspring who had increased concentrations compared to HSHFD fed control males (Fig 3H-I). In addition, the reduction in circulating triglycerides was significant for HSHFD fed UE, but not in control female offspring. Overall, these data demonstrate that placental endocrine malfunction, via modification of an imprinting gene, induces programming effects in the adult offspring. Namely, both UE female and male offspring are programmed for insulin resistance, and are predisposed to a greater gain of adiposity on an obesogenic diet.

## Discussion

From an evolutionary point of view, insulin resistance during pregnancy is designed to limit maternal glucose utilization by target tissues, and thereby shunt an adequate amount of supply to the developing fetus. In humans, there is a linear relationship between maternal glucose tolerance during pregnancy and birthweight (HAPO Study Cooperative Research Group et al., 2008). The mechanisms that render the mother’s tissues less insulin sensitive are poorly understood, but are believed to be caused, at least in part, by placental hormones (Napso et al., 2018; Stern et al., 2021). To our knowledge, the role of the placenta, or of a single hormone, in this process has not been directly demonstrated. Here, we have shown in the mouse that IGF2, a peptide with structural similarities to insulin, produced in placental endocrine cells is required to establish the insulin resistance state in pregnancy.

Our data indicate that loss of *Igf2* from the endocrine layer of the mouse placenta likely alters the secretion of key signalling proteins into the maternal circulation, which in turn, leads to an impaired maternal endocrine and lipid profile. Ultimately, these events render maternal organs more insulin sensitive, with important consequences for fetal development. Our data strongly suggests that loss of fetal viability and the fetal growth restriction are caused by insufficient glucose and lipid delivery to the fetus, mainly driven by the higher glucose utilization/uptake by the maternal organs as a result of increased glucose tolerance and insulin sensitivity.

Another major finding of our work is that the pancreatic β-cell mass in mothers that carry *Igf2* mutant UE placentas fail to expand compared to pregnant controls. Pancreatic β-cell mass expansion is thought to occur in response to increased insulin resistance in maternal tissues (Rieck and Kaestner, 2010). Our findings may therefore reflect the failure in acquiring the insulin resistance state. We suggest that *Igf2* mutant UE placentas fail to secrete hormones that are necessary to stimulate β-cell expansion, such as prolactins, which are decreased in circulation of UE dams. Prior work using litters comprised of both control and UE fetuses has reported that *Igf2* deficiency in placental endocrine cells indeed reduces the expression of prolactin genes, and disrupts the expression of *Cyp17a1,* which would favour the production of estrogen over progesterone by the placenta (Aykroyd et al., 2020). Herein, we found altered secretion or circulating levels of previously described regulators of insulin sensitivity during gestation, namely estradiol/progesterone ratio, prolactins and TNFA. As novel mediators, we found reductions in circulating levels of serine protease inhibitors, including kallistatin. In the liver, the observed increases in PPARs, mTOR and LPL abundance may contribute to the reduction in circulating lipids, and the tendency for decreased PEPCK would suggest inhibition of glucose production by the liver (another major maternal metabolic adaptation that ensures glucose delivery to the fetus).

*Igf2* imprinted expression in placenta endocrine cells may have evolved as a strategy to mobilize nutrients to the growing fetus, in line with the concept of imprinted genes having evolved due to conflict (Haig, 1993). Based on our targeted loss-of-function results, we suggest that placental *Igf2* imprinting is a major modulator of maternal metabolic adaptations that ensures mobilization of nutrients, namely glucose and lipids, to the fetus by inducing insulin resistance and enhancing maternal glucose and lipid availability. This proposal is further supported by circumstantial evidence collected in mice and human studies (Jacob et al., 2013), including from human imprinted disorders Beckwith-Wiedemann Syndrome (BWS), an overgrowth syndrome, and Silver-Russell Syndrome (SRS), a fetal growth restriction syndrome. In a mouse model of BWS, mothers that carry overgrown offspring with increased *Igf2* and reduced levels of the maternally expressed *H19* gene in most fetal cells develop maternal hyperglycemia and glucose intolerance (Petry et al., 2010). Metabolic phenotypes in offspring such as neonatal hyperinsulinemia and hypoglycaemia have also been reported in subgroups of BWS and SRS patients, which show high *IGF2*/low *H19* and low *IGF2*/increased *H19* expression levels in tissues (Sparago et al., 2004; Yamazawa et al., 2008), respectively.

Currently, there are no data to support a role for placental imprinting in fetal programming of metabolic disease, mainly due to the lack of appropriate, specific genetic tools. We found that male and female offspring with *Igf2* deficiency in placental endocrine cells become insulin resistant and have elevated adiposity, with this effect stronger particularly under conditions that favour the mismatch of programmed adaptations *in utero* (i.e. intrauterine growth restriction due to compromised nutrient supply, with a postnatal obesogenic environment). We conclude that placental endocrine malfunction via manipulation of *Igf2* in the placental endocrine cells programmes obesity and insulin resistance in the adult offspring.

In conclusion, this study provides eagerly-awaited experimental evidence for an intrinsic fetal manipulation system, which operates in the placenta to modify maternal metabolism and nutrient partitioning to the fetus, with implications for the metabolic health and disease risk of offspring in later life.

## Materials and Methods

### Mice and genotyping

Mice were housed at the University of Cambridge Animal Facility under a 12/12 dark/light system. All experiments were performed under the U.K. Animals (Scientific Procedures) Act 1986, after ethical approval by the University of Cambridge. Both *Igf2^Flox^*and *Tpbpa^Cre^* mouse lines have been on the C57BL/6J genetic background for more than 10 generations (Hammerle et al., 2020; Simmons et al., 2007)^1,2^. The genotype of mouse lines was confirmed by PCR analysis of genomic DNA extracted from ear biopsies. Details of the primer sequences for genotyping and about the generation of both genetic lines have been reported previously (Aykroyd et al., 2020). To visualize the Cre recombinase activity, *Tpbpa^Cre^* mice were mated with *Rosa26*TdRFP reporter mouse line [*Gt(ROSA)26Sor;* Jackson lab and was generously provided by Prof William Colledge at the University of Cambridge] (Luche et al., 2007).

For studies of pregnancy physiology and fetal outcomes, virgin female (♀) homozygous *Tpbpa^Cre^* mice were mated with male (♂) homozygous *Igf2^Flox^* to generate pregnancies where the entire litter had selective loss of *Igf2* in the endocrine cells of the placental junctional zone. The opposite parental mating strategy (virgin ♀ homozygous *Igf2^Flox^* mated with ♂ homozygous *Tpbpa^Cre^*) generated pregnancies with control litters. The day of the copulatory plug was defined as day 1 of pregnancy (GD1) and all procedures related to pregnant females were performed on GD16. Age-matched virgin ♀ homozygous *Tpbpa^Cre^* and ♀ homozygous *Igf2^Flox^* were used for pre-pregnancy baseline values.

For offspring studies, dams were allowed to deliver naturally and litters were standardised to six pups on postnatal day 3 (3 males and 3 females, whenever possible). Pups were weaned on day 21 and, thereafter, female and male offspring were weighed weekly.

At 12 weeks of age, one animal per sex and litter received an intraperitoneal glucose or an insulin tolerance test. All offspring were euthanized at 13 weeks of age, and one animal per sex and litter underwent body composition analysis using DEXA scanning on a Lunar PIXImus densitometer (GE Lunar Corp., Madison, WI, USA).

### Diets

Two different diets were used for this study. Pregnant dams were fed *ad libitum* throughout pregnancy and lactation with a standard chow RM3 diet obtained from Special Dietary Services (Witham, UK). For the offspring studies, entire litters were randomised immediately after weaning to either the standard RM3 chow, or an obesogenic diet with a 45% kcal fat (#D12451, Research Diets, New Brunswick, NJ, USA) and 20% sucrose in tap water supplemented with AIN93G mineral and vitamin mix (MP Biomedicals, Irvine, CA, USA).

### Glucose and insulin tolerance tests

Mice were fasted for 6 hours from 08:00h on the test day. A blood sample and glucose measure was obtained from a small cut in the tail vein immediately before the intraperitoneal injection of glucose (10% weight for volume, 1 g/kg body weight) or insulin (0.75 units/kg, Actrapid; Novo Nordisk, Denmark). Blood glucose levels were quantified at 0, 15, 30, 60, 90, and 120LJminutes post injection.

### Blood collection and metabolite and hormone concentrations

Mice were anesthetised with fentanyl-fluanisone and midazolam in sterile water (10µl/g mouse, 1:1:2 ratio intraperitoneally), and blood was collected by cardiac exsanguination followed by cervical dislocation. Blood glucose was measured using a handheld glucometer (One Touch Ultra; LifeScan Inc., Malvern, PA, USA). Blood samples in EDTA-coated tubes were centrifuged at 1,500 rpm, plasma collected and stored at -20°C. Enzymatic assay kits were used to measure plasma insulin (Mercodia: 10-1247-01, Uppsala, Sweden), estradiol (Cayman Chemical: 582251, Ann Arbor, MI, USA), progesterone (Cayman Chemical: 582601), IGF2 (R&D Systems: DY792, Minneapolis, MN, USA), NEFAs (Sigma-Aldrich Corp: 11383175001, St Louis, MO, USA), cholesterol (Siemens Healthcare: DF27, Erlangen, Germany), and triglycerides (Siemens Healthcare: DF69A) as per manufacturer instructions.

### Pancreas insulin content and pancreas immunostaining

Maternal pancreas was removed, weighed and either snap-frozen in liquid nitrogen for biochemical analysis or fixed in 4% paraformaldehyde for paraffin embedding and histological assessment. For pancreas insulin content, whole, snap-frozen pancreas were crushed, mixed with ethanol/hydrochloric acid, and incubated overnight at -20°C. The tissue was then disrupted using a Dounce homogeniser, left overnight at 4°C and subsequently centrifuged. The supernatant was taken to measure insulin as described above. Values were expressed per mg of tissue. For insulin staining, portions of the entire pancreas were sectioned at 5μm thickness and ≥3 non-consecutive sections of each portion were stained with an antibody against insulin (1:100, Cell Signaling Technologies: 4590, Danvers, MA, USA), which was subsequently detected by biotinylated goat anti-rabbit secondary antibody (1:1000, Abcam: ab6720, Cambridge, UK), and visualised using streptavidin-conjugated peroxidase (1:500, Rockland Immunochemicals: S000-03, Gilbertsville, PA, USA) and 3,3′-Diaminobenzidine (DAB, Abcam: ab64238). Slides were counterstained with nuclear fast red (Vector Laboratories: H-3403, Burlingame, CA, USA) and sections were digitalized using a nanozoomer scanner (Hamamatsu Photonics, Shizuoka Prefecture, Japan). Analysis was performed using Image J software (National Institutes of Health, Bethesda, MD, USA) and conducted blinded to experimental groups.

### Placental IGF2 immunostaining

Paraffin-embedded placental sections cut at 5 µm were used. Antigen retrieval was performed with in 10mM sodium citrate. Slides were incubated overnight with goat anti-human IGF-II antibody (R&D Systems; AF292). Alexa Fluor® 488-AffiniPure antibody (Jackson ImmunoResearch; 705-546-147) and DAPI (Sigma; D9542) were used to visualize IGF2 immuno-staining and nuclei in the section, respectively. Image acquisition was performed with a LSM510 Meta confocal laser scanning microscope (Carl Zeiss, Jena, Germany) and ZEN 2009 software.

### Fetal growth and placental histological analysis

A representative number of litters were kept to assess fetal symmetry. In particular, fetuses were photographed using a Zeiss SteREO Discovery V8 microscope with an AxioCam MRc5 camera and imaged with AxioVision 4.7.2 software (Carl Zeiss AG, Oberkochen, Germany). Biometric measurements were performed using Image J software. Placentas were fixed, paraffin-embedded, exhaustively sectioned at 8μm and stained with haematoxylin and eosin to analyse junctional zone volume and endocrine cell populations using the Computer Assisted Stereological Toolbox (CAST v2.0) program as previously described^3^. Analyses were performed blinded to the genotype.

### Placental qRT-PCR

RNA was extracted from micro-dissected placental layers performed as previously described (Aykroyd et al., 2020). Following cDNA synthesis with Applied Biosystems High-Capacity cDNA Reverse Transcription Kit (Thermo Fisher Scientific, Waltham, MA, USA), qRT-PCR was performed using the primers sequences specified in Supplementary Table 3. Relative mRNA expression levels were then calculated using the 2^-ΔΔCT^ method (Livak and Schmittgen, 2001).

### Placental glucose transport assay

To assess unidirectional materno-fetal clearance of the non-metabolisable radioactive tracer ^3^H-methyl D-Glucose (MeG), pregnant dams mice were anaesthetised and injected intravenously as previously described (Sferruzzi-Perri, 2018; López-Tello et al., 2019). Fetuses were then decapitated and lysed at 55°C in Biosol (National Diagnostics, Atlanta, GA USA). Fetal lysates were measured for beta emissions by liquid scintillation counting (Optiphase Hisafe II and Packard Tri-Carb, 1900; Perkin-Elmer, Waltham, MA, USA) and radioactivity (DPM; disintegrations per minute).

### Primary cell cultures of mouse placenta cells and LC-MS/MS analysis of conditioned media

All placentas within representative litters per genotype were pooled and primary placental endocrine cell cultures were performed as described previously (Napso et al., 2021). Briefly, placentas were enzymatically dissociated and grown in a humidified atmosphere of 5% CO_2_ at 37°C. Cells were maintained with NTCT-135 medium containing 10% fetal bovine serum, 50 IU/ml ampicillin, 50 µg/ml streptomycin, and 2mM l-glutamine. Cell medium was replaced at 24h and the final length of culture was 48h. Conditioned medium from cultured placental cells was collected at 48h of culture and subjected to LC-MS/MS analysis. Of note, 24h prior to the collection of the conditioned medium, cells were washed three times with PBS and cultured in serum-free medium. The conditioned media was centrifuged for 10 min at 1000g. All LC-MS/MS experiments and data analysis has been described elsewhere^4^. In particular, UniProt Knowledgebase (UniProtKB) gene ontology analysis (https://www.uniprot.org/) was performed to identify relev ant pathways that proteins uniquely present in the control and UE secretomes (165 and 125, respectively) are involved in.

### LC-MS/MS analysis of mouse plasma

Extraction and analysis of circulating proteins in plasma was performed as described elsewhere (Napso et al., 2021). Briefly, mouse plasma was defrosted and 6M guanidine hydrochloride was added in 1:1 proportion. Precipitation of proteins was performed by adding 75% acetonitrile in water at a ratio of 1:6 (diluted plasma:acetonitrile) followed by sample centrifugation at 2900 × *g* for 10 min at 4°C. Extracts were acidified and analysed using nano LC-MS/MS and the data searched using PEAKS. Peptides from specific proteins of interest in the PEAKS search result were selected for targeted quantitation in the raw LC-MS/MS files. The peak area of each peptide was expressed as a ratio of a digested peptide from the spiked internal standard peptide (bovine insulin).

### Western immunoblotting

Protein was extracted from maternal liver using RIPA buffer (Sigma-Aldrich Corp., R0278). Protein concentration was determined by a Pierce™ BCA Protein Assay Kit (Thermo Fisher Scientific). After electrophoresis and protein transfer onto 0.2μm nitrocellulose membranes (Bio-Rad Laboratories Inc., Hercules, CA, USA), membranes were blocked in fetal bovine serum or skim milk and incubated with primary antibodies (Supplementary Table 4) overnight on a rocker at 4°C. Following washes, membranes were incubated with secondary antibody (dilution 1:10,000; NA931 or NA934; Amersham ECL Mouse/Rabbit IgG, HRP-linked). The iBright 1500 Imaging System (Invitrogen) was used to detect immunocomplexes in the membrane after application of SuperSignal™ West Femto Maximum Sensitivity Substrate (Thermo Fisher Scientific). ImageJ software (1.49version) was used to quantitate the densitometric intensity of immunoreactive bands. To check for even protein loading, membranes were stained and normalized with Ponceau S (Romero-Calvo et al., 2010).

### Statistical analysis

All data are shown with individual data points and the mean±SEM where possible. Feto-placental weights and placental transport assays were analysed by ANOVA with repeated measures (MIXED model) using the SAS/STAT Software (Statistical System Institute Inc. Cary, NC, USA). Each mother was considered the subject, the genotype the main factor and litter size was added as covariate. Offspring biometry data were analysed using litter means by two-way ANOVA. Glucose and insulin tolerance tests were analysed by repeated measures ANOVA followed by Tukey post-hoc test for mean comparisons. Maternal data were analysed by ANOVA using GraphPad Prism software (GraphPad Software Inc., La Jolla, CA, USA). For analyses only using two groups (namely litter size, steroid concentrations, and western blotting data), the Shapiro-Wilk test was performed to determine if data followed the normal distribution and then statistical analysis was performed using Mann Whitney or unpaired student’s t-tests, as required. The numbers of samples used for each experiment are shown within the respective figure or figure legends. A value of p<0.05 was considered statistically significant unless stated otherwise.

## Data availability

The authors declare that the data generated and analysed to support the findings of this study are available within the paper and its supplementary files, or available from the authors upon reasonable request.

## Acknowledgments

We would like to thank Dr Richard Kay and Dr Amy George for LC-MS/MS analysis (Wellcome-MRC Institute of Metabolic Science); Dr Keith Burling lab (Biochemical Assay Laboratory) for performing hormone/metabolites quantification and Mrs Marta Ibañez for bioinformatics assistance.

## Funding

JL-T currently holds a Sir Henry Wellcome Postdoctoral Fellowship from the Wellcome Trust [Grant number 220456/Z/20/Z] and previously a Newton International Fellowship from the Royal Society [NF170988 / RG90199] and an Early Career Grant from the Society for Endocrinology. HEJY was supported by an A*STAR International Fellowship from the Agency for Science, Technology and Research. ESP was supported by a Beca-Chile, ANID Postdoctoral Scholarship: 74190055. ERC was supported by a Cambridge-Rutherford PhD Scholarship from the Cambridge Trust and Rutherford Foundation. TN was supported by an EU Marie Skłodowska-Curie Fellowship (PlaEndo/703160) and an Early Career Grant from the Society for Endocrinology. ANSP was supported by a Royal Society Dorothy Hodgkin Research Fellowship, Academy of Medical of Sciences Springboard Grant, Isaac Newton Trust Grant and Lister Institute Research Prize grant to ANSP (grant numbers: DH130036 / RG74249, SBF002/1028 / RG88501, RG97390 and RG93692, respectively). M.C. research was supported by the Biotechnology and Biological Sciences Research Council (grant BB/H003312/1) and the Medical Research Council (MRC_MC_UU_12012/4 to M.C.; MRC_MC_UU_12012/5 to the MRC Metabolic Diseases Unit).

## Competing interests

The authors declare no competing interests

---

**Supplementary Figure 1.**
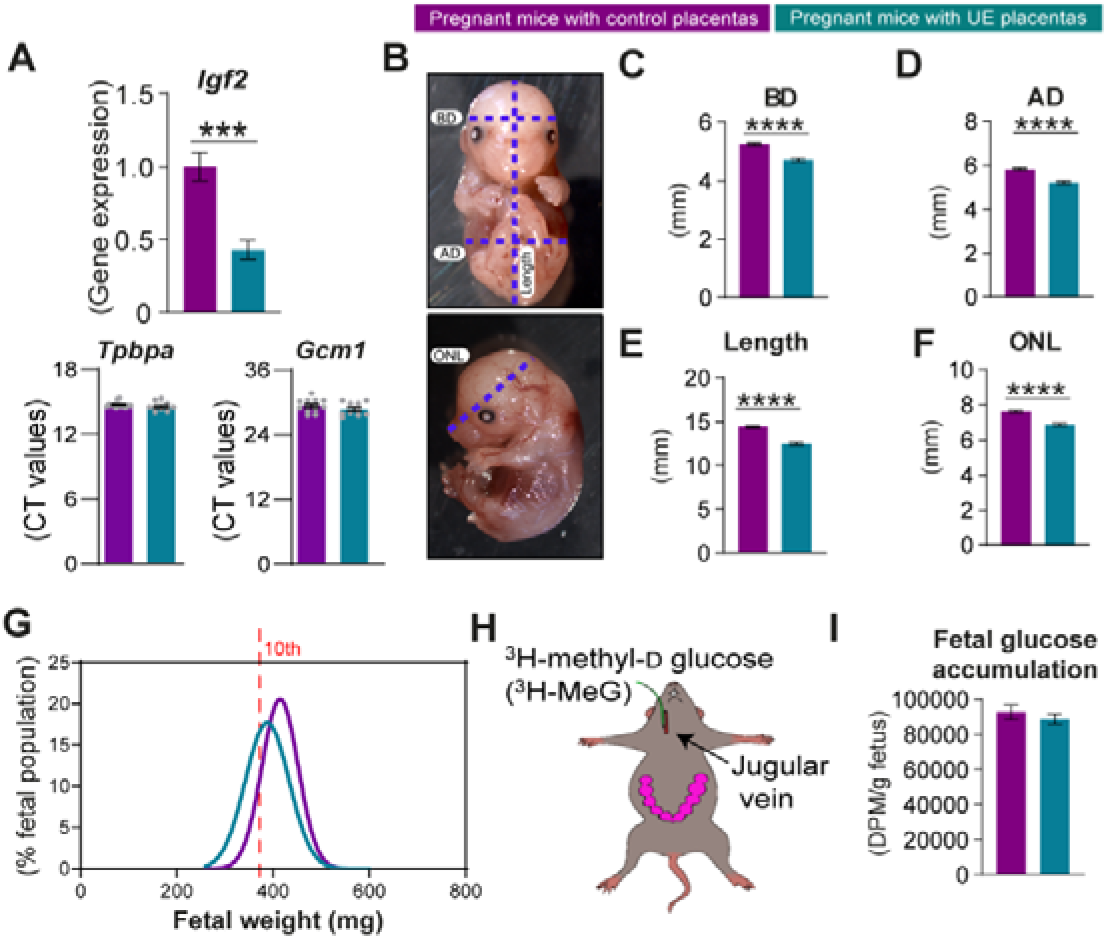
Additional fetal growth parameters and fetal glucose accumulation at GD16. (A) Gene expression levels of *Igf2*, and cycle threshold (CT) values of *Tpbpa* (junctional zone marker) and *Gcm1* (labyrinth zone marker) in dissected junctional zone samples that have some labyrinth zone contamination (n=10-13 placentas/group). (B) Representative image showing measured fetal growth parameters of biparietal diameter (BD), abdominal diameter (AD), length and occipital to nose length (OCL). (C-F) Measurements of fetal growth parameters (n=6 litters/group). (G) Distribution of fetal weights with line showing 10^th^ centile cut-off (372mg, representing 10^th^ centile of control fetuses). (H) Schematic of jugular vein injection with radio-labelled glucose for placental transport assay. (I) Fetal glucose accumulation following injection of dam with radio-labelled glucose (n=7-8 litters/group). Data presented as mean±SEM with or without individual data points shown. Fetal data were analysed by ANOVA with repeated measures (MIXED model), considering the mother as the subject and the genotype the main factor, with litter size added as a covariate. ****p<0.0001.

**Supplementary Figure 2.**
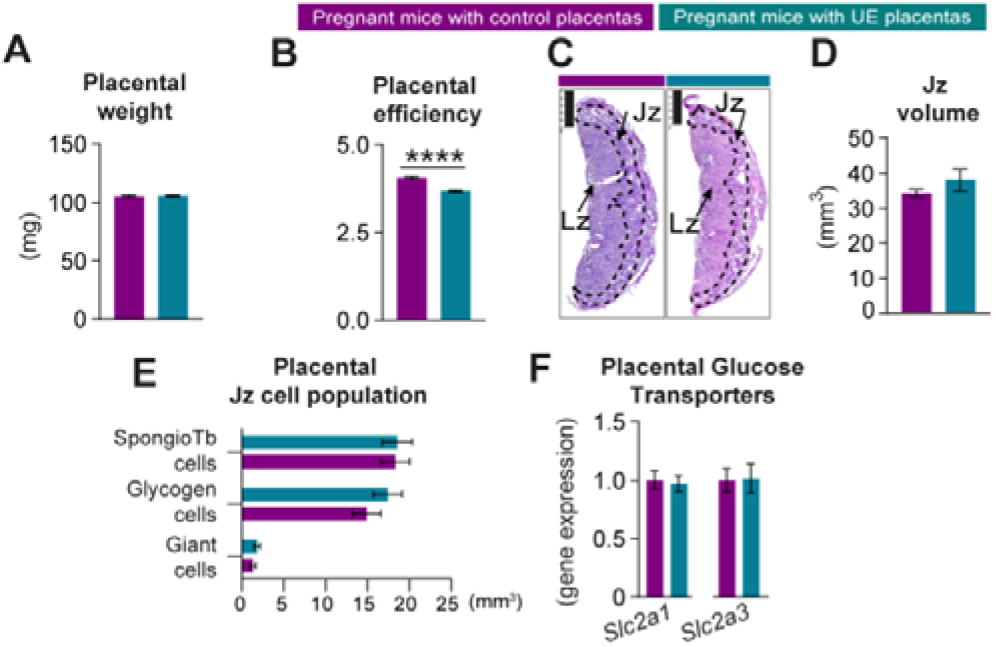
Placental parameters at GD16. (A) Placental weight (n=28-32 litters/group). (B) Placental efficiency defined by ratio of fetal weight to placental weight (n=28-32 litters/group). (C) Representative histological sections of control and UE placentas with marked junctional zone (Jz) and labyrinthine zone (Lz) regions. Scale bar is 1mm. (D) Junctional zone volume as determined by placental stereological analysis (n=5/group). (E) Proportions of spongiotrophoblast, glycogen and giant cell populations within the junctional zone (n=4-5/group). (F) Relative mRNA expression of placental glucose transporters (n=14 fetuses/6 litters/group) Data presented as mean±SEM. Feto-placental data were analysed by ANOVA with repeated measures (MIXED model), considering the mother as the subject and the genotype the main factor, with litter size added as a covariate. ****p<0.0001.

**Supplementary Figure 3.**
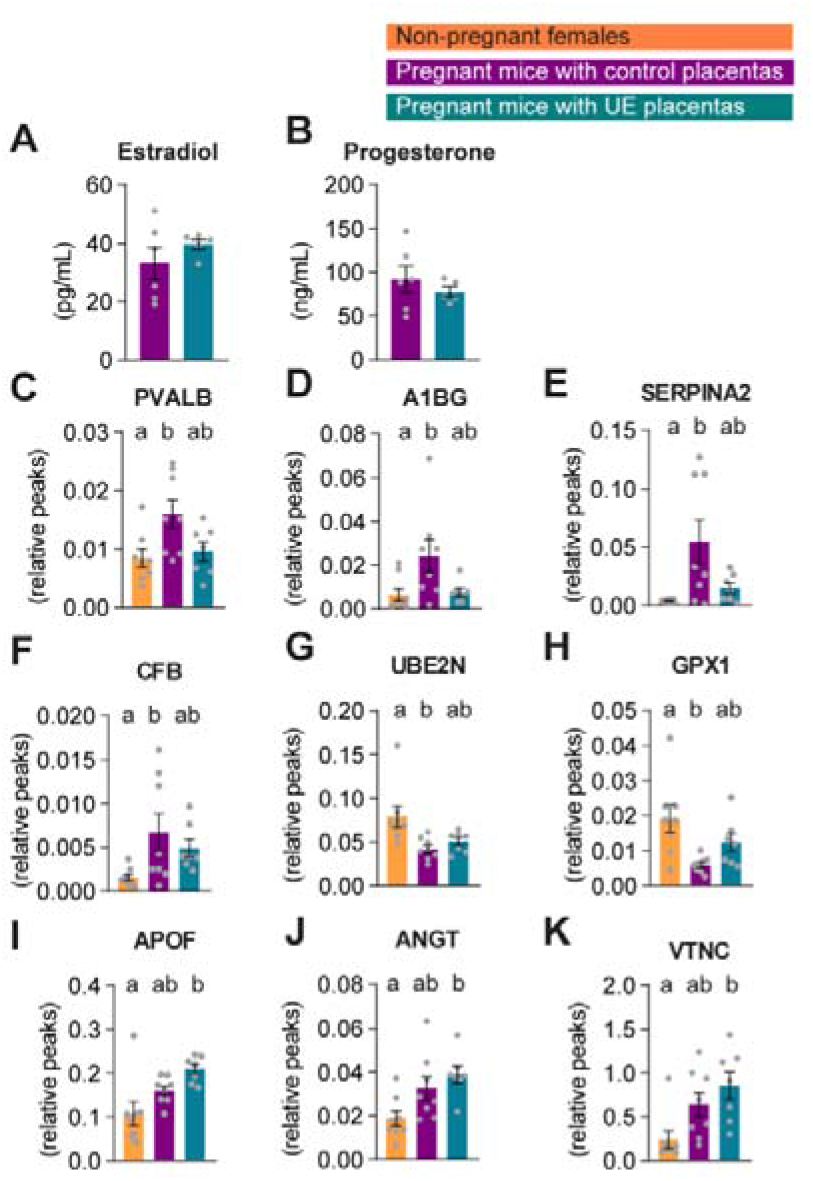
Measurements of circulating steroid hormones and proteins in maternal blood at GD16 and non-pregnant females. (A-B) Circulating concentrations of steroid hormones, estradiol and progesterone as determined by ELISA (n=5-6/group). (C-K) Relative plasma levels of proteins identified by LC-MS/MS in the circulation of UE dams that were intermediate between non-pregnant and pregnant state (C-H) or significantly different to non-pregnant state (I-K) (n=7-8/group). Data presented as mean±SEM with individual data points shown. ANOVA was performed with Tukey post-hoc tests for mean comparisons. Different letters (a, b) indicate significant differences between groups by ANOVA (p<0.05).

**Supplementary Figure 4.**
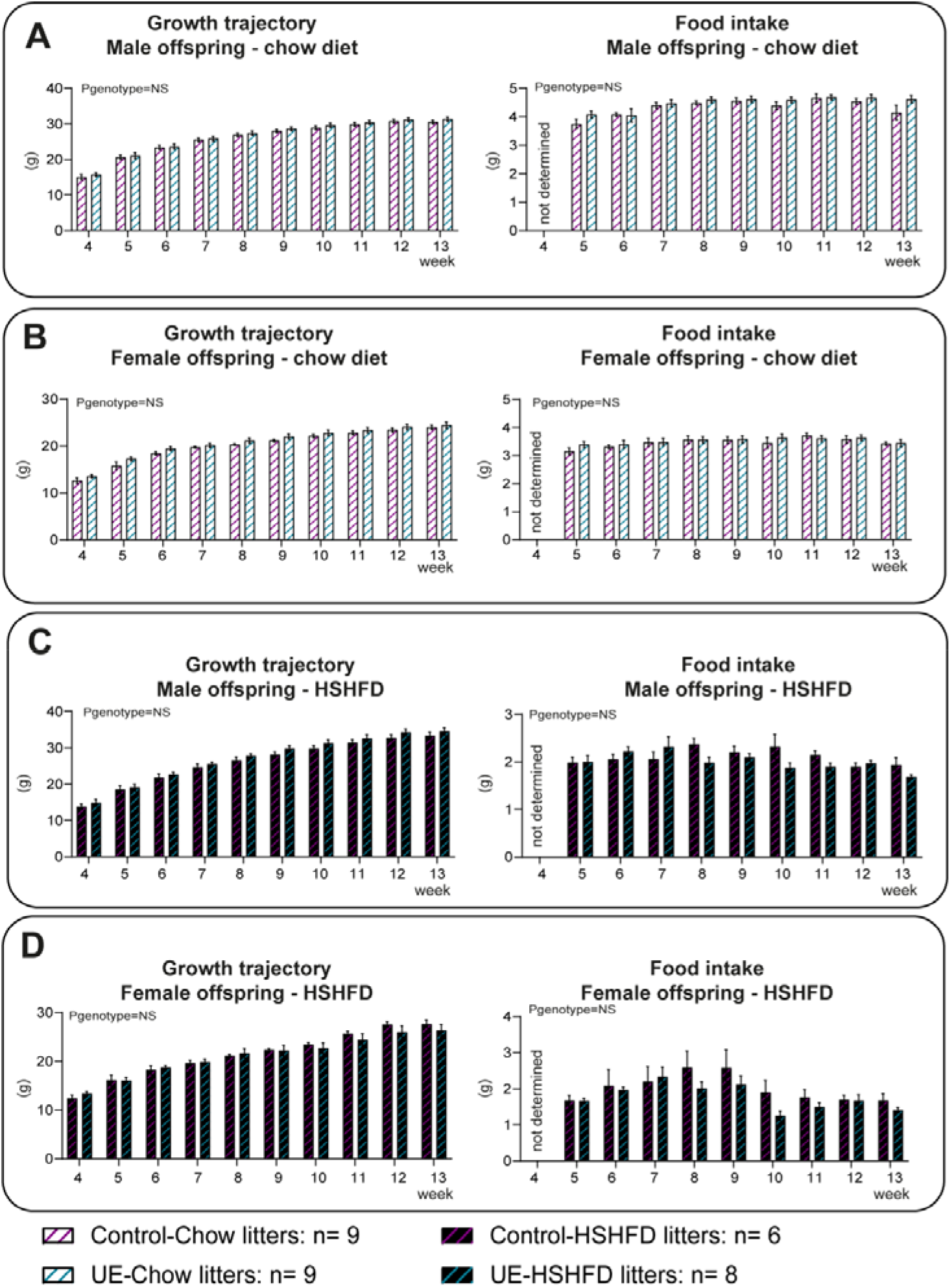
Growth trajectories and food intake of male and female offspring. (A-D) Weekly measures of weight and food intake of male and female offspring supported by control and UE placentas from 4 weeks and 5 weeks of age respectively. (n=6-9/group) Data presented as mean±SEM. Two-ways ANOVA repeated measures with Bonferroni post-hoc tests for mean comparisons.

**Supplementary Figure 5.**
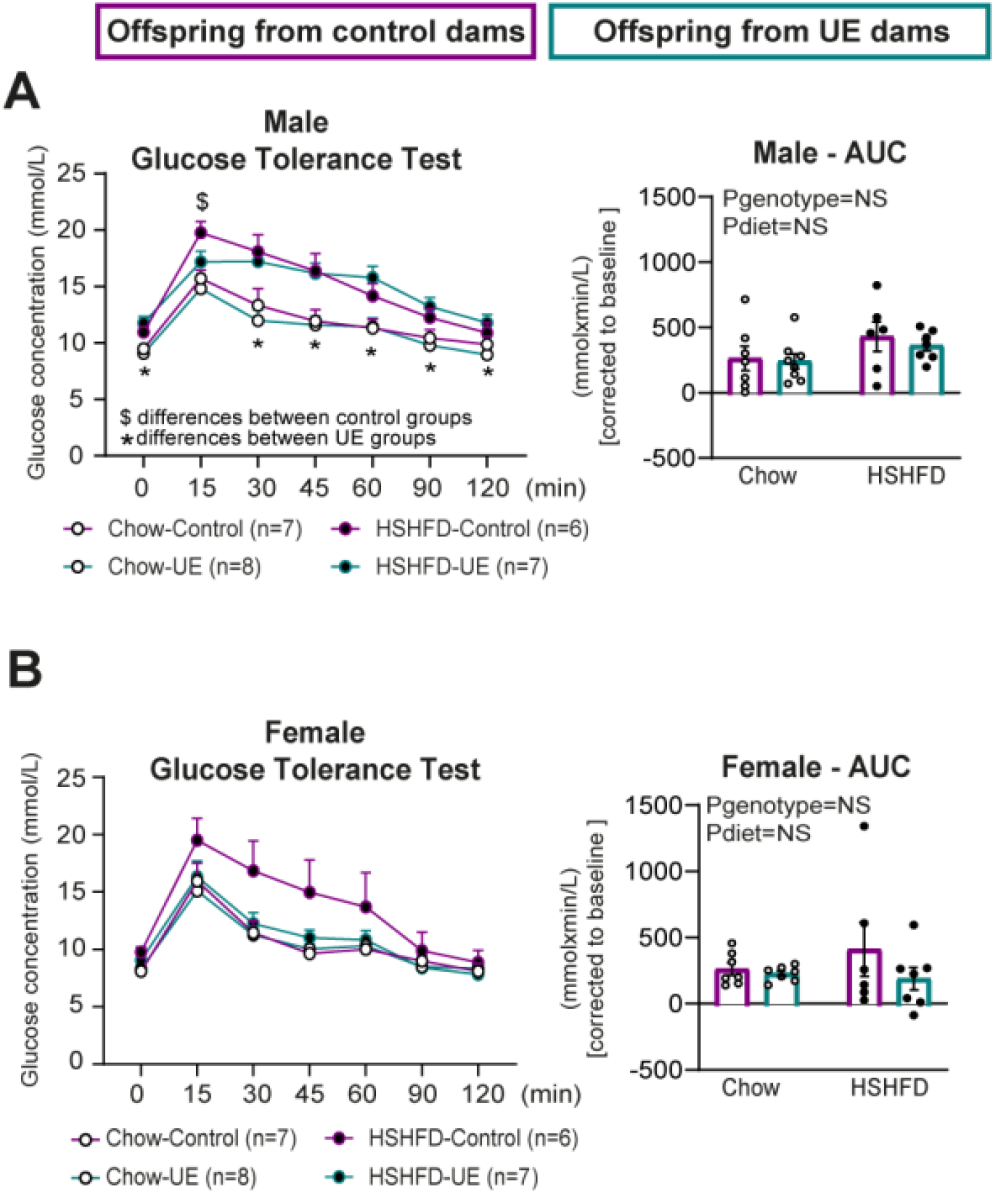
Glucose tolerance tests of male and female offspring on chow and high sugar high fat diet (HSHFD) (A-B) Intraperitoneal glucose tolerance tests were performed at 12 weeks of age. Inserts show area under glucose curve (AUC) (n=6-8/group/sex). Data presented as mean±SEM with individual data points shown. Two-ways ANOVA was performed with Tukey post-hoc test for mean comparisons. Different symbols (*, $) indicate significant differences between groups by two-ways ANOVA repeated measures.

**Supplementary Table 1. List of proteins and Uniprot analysis of proteins detected in the placental secretome by liquid chromatography mass spectrometry (LC-MS/MS).**

**Supplementary Table 2. List of proteins detected in the maternal plasma by liquid chromatography mass spectrometry (LC-MS/MS).**

**Supplementary Table 3.**
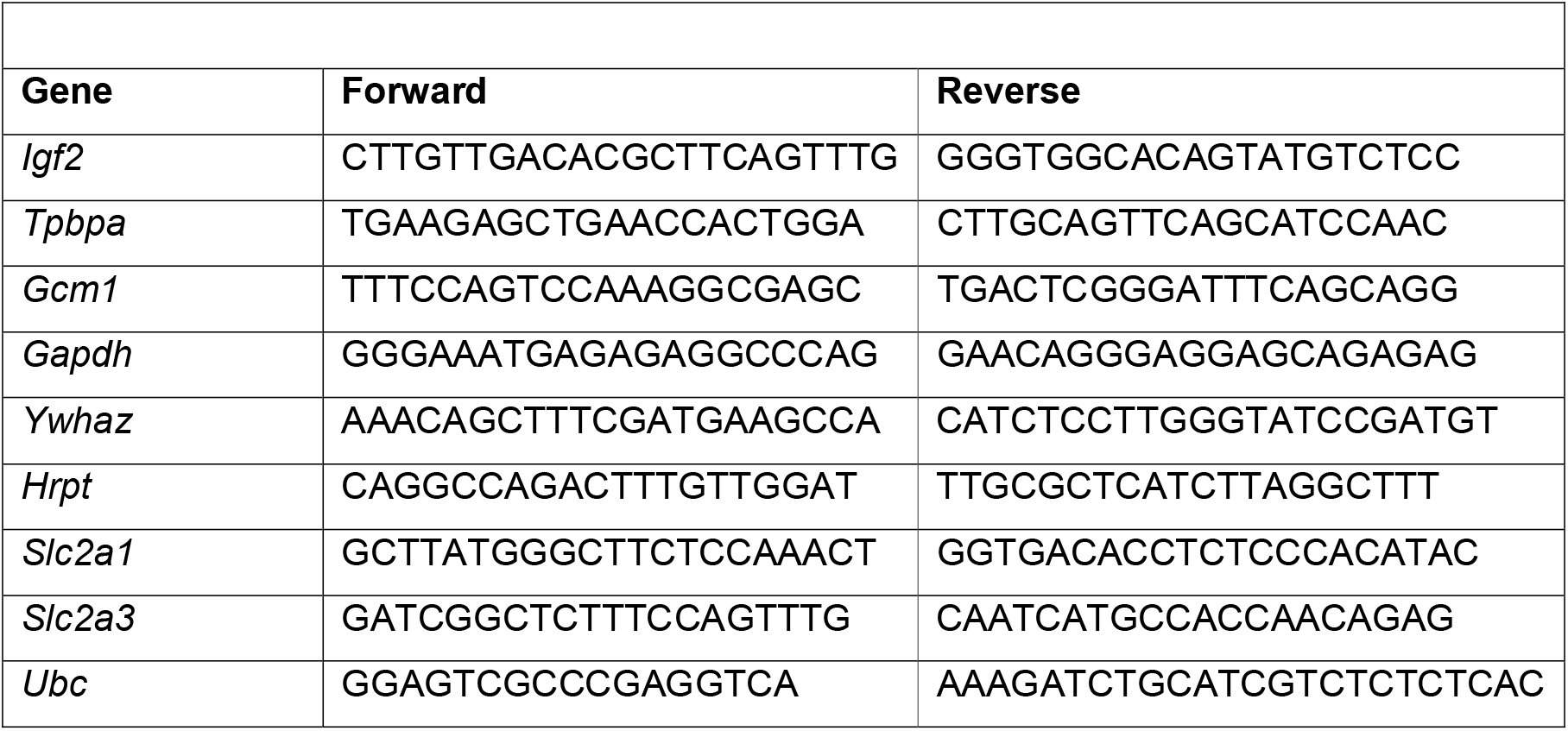
List of primers used for qPCR.

**Supplementary Table 4.** List of antibodies used for western blotting.

| Supplementary Table 4 |  |  |
| --- | --- | --- |
| Antibody | Catalogue number | Dilution |
| Insulin receptor beta | Cell Signalling, 3020 | 1:1000 |
| PI3K-p85 alpha | Millipore, 06-195 | 1:5000 |
| Phospho-Akt (Ser473) | Cell Signalling, 4060 | 1:1000 |
| AKT | Cell Signalling, 9272 | 1:1000 |
| GβL (86B8) | Cell Signalling, 3272 | 1:1000 |
| PEPCK | Santa Cruz, sc-271029 | 1:200 |
| Lipoprotein lipase | Abcam, ab21356 | 1:1000 |
| PPAR alpha | Abcam, ab3484 | 1:1000 |
| PPAR gamma | Santa Cruz, sc-7273 | 1:1000 |

## Notes

### Competing Interest Statement

The authors have declared no competing interest.

## References

Akash, M.S.H., Rehman, K., and Liaqat, A. (2018). Tumor Necrosis Factor-Alpha: Role in Development of Insulin Resistance and Pathogenesis of Type 2 Diabetes Mellitus. J Cell Biochem 119, 105–110. https://doi.org/10.1002/jcb.26174.

Aykroyd, B.R.L., Tunster, S.J., and Sferruzzi-Perri, A.N. (2020). Igf2 deletion alters mouse placenta endocrine capacity in a sexually dimorphic manner. J. Endocrinol. 246, 93–108. https://doi.org/10.1530/JOE-20-0128.

Baeyens, L., Hindi, S., Sorenson, R.L., and German, M.S. (2016). β-Cell adaptation in pregnancy. Diabetes Obes Metab 18 *Suppl 1*, 63–70. https://doi.org/10.1111/dom.12716.

Berthier, A., Johanns, M., Zummo, F.P., Lefebvre, P., and Staels, B. (2021). PPARs in liver physiology. Biochim Biophys Acta Mol Basis Dis 1867, 166097. https://doi.org/10.1016/j.bbadis.2021.166097.

Borges, B.C., Han, X., Allen, S.J., Garcia-Galiano, D., and Elias, C.F. (2019). Insulin signaling in LepR cells modulates fat and glucose homeostasis independent of leptin. Am J Physiol Endocrinol Metab 316, E121–E134. https://doi.org/10.1152/ajpendo.00287.2018.

Cao, Y., Li, X., Lu, C., and Zhan, X. (2018). The relationship between vitronectin and hepatic insulin resistance in type 2 diabetes mellitus. Endocr J 65, 747–753. https://doi.org/10.1507/endocrj.EJ17-0504.

Cassidy, F.C., and Charalambous, M. (2018). Genomic imprinting, growth and maternal-fetal interactions. J Exp Biol 221, jeb164517. https://doi.org/10.1242/jeb.164517.

Coan, P.M., Barrier, M., Alfazema, N., Carter, R.N., Marion de Procé, S., Dopico, X.C., Garcia Diaz, A., Thomson, A., Jackson-Jones, L.H., Moyon, B., et al. (2017). Complement Factor B Is a Determinant of Both Metabolic and Cardiovascular Features of Metabolic Syndrome. Hypertension HYPERTENSIONAHA.117.09242. https://doi.org/10.1161/HYPERTENSIONAHA.117.09242.

Constância, M., Kelsey, G., and Reik, W. (2004). Resourceful imprinting. Nature 432, 53–57. https://doi.org/10.1038/432053a.

DeChiara, T.M., Robertson, E.J., and Efstratiadis, A. (1991). Parental imprinting of the mouse insulin-like growth factor II gene. Cell 64, 849–859. https://doi.org/10.1016/0092-8674(91)90513-x.

Essaji, Y., Yang, Y., Albert, C.J., Ford, D.A., and Brown, R.J. (2013). Hydrolysis products generated by lipoprotein lipase and endothelial lipase influence macrophage cell signalling pathways. Lipids 48, 769–778. https://doi.org/10.1007/s11745-013-3810-6.

Fowden, A.L., and Moore, T. (2012). Maternal-fetal resource allocation: co-operation and conflict. Placenta 33 *Suppl 2*, e11–15. https://doi.org/10.1016/j.placenta.2012.05.002.

Geng, L., Lam, K.S.L., and Xu, A. (2020). The therapeutic potential of FGF21 in metabolic diseases: from bench to clinic. Nat Rev Endocrinol 16, 654–667. https://doi.org/10.1038/s41574-020-0386-0.

Haig, D. (1993). Genetic conflicts in human pregnancy. Q Rev Biol 68, 495–532. https://doi.org/10.1086/418300.

Hammerle, C.M., Sandovici, I., Brierley, G.V., Smith, N.M., Zimmer, W.E., Zvetkova, I., Prosser, H.M., Sekita, Y., Lam, B.Y.H., Ma, M., et al. (2020). Mesenchyme-derived IGF2 is a major paracrine regulator of pancreatic growth and function. PLOS Genetics 16, e1009069. https://doi.org/10.1371/journal.pgen.1009069.

HAPO Study Cooperative Research Group, Metzger, B.E., Lowe, L.P., Dyer, A.R., Trimble, E.R., Chaovarindr, U., Coustan, D.R., Hadden, D.R., McCance, D.R., Hod, M., et al. (2008). Hyperglycemia and adverse pregnancy outcomes. N Engl J Med 358, 1991–2002. https://doi.org/10.1056/NEJMoa0707943.

Jacob, K.J., Robinson, W.P., and Lefebvre, L. (2013). Beckwith-Wiedemann and Silver-Russell syndromes: opposite developmental imbalances in imprinted regulators of placental function and embryonic growth. Clin Genet 84, 326–334. https://doi.org/10.1111/cge.12143.

Jenkins, A.J., McBride, J.D., Januszewski, A.S., Karschimkus, C.S., Zhang, B., O’Neal, D.N., Nelson, C.L., Chung, J.S., Harper, C.A., Lyons, T.J., et al. (2010). Increased serum kallistatin levels in type 1 diabetes patients with vascular complications. J Angiogenes Res 2, 19. https://doi.org/10.1186/2040-2384-2-19.

Killian, J.K., Nolan, C.M., Stewart, N., Munday, B.L., Andersen, N.A., Nicol, S., and Jirtle, R.L. (2001). Monotreme IGF2 expression and ancestral origin of genomic imprinting. J Exp Zool 291, 205–212. https://doi.org/10.1002/jez.1070.

Kim, D.-H., Sarbassov, D.D., Ali, S.M., Latek, R.R., Guntur, K.V.P., Erdjument-Bromage, H., Tempst, P., and Sabatini, D.M. (2003). GbetaL, a positive regulator of the rapamycin-sensitive pathway required for the nutrient-sensitive interaction between raptor and mTOR. Mol Cell 11, 895–904. https://doi.org/10.1016/s1097-2765(03)00114-x.

Kolodziejski, P.A., Sassek, M., Bien, J., Leciejewska, N., Szczepankiewicz, D., Szczepaniak, B., Wojciechowska, M., Nogowski, L., Nowak, K.W., Strowski, M.Z., et al. (2020). FGF-1 modulates pancreatic β-cell functions/metabolism: An in vitro study. Gen Comp Endocrinol 294, 113498. https://doi.org/10.1016/j.ygcen.2020.113498.

Livak, K.J., and Schmittgen, T.D. (2001). Analysis of relative gene expression data using real-time quantitative PCR and the 2(-Delta Delta C(T)) Method. Methods 25, 402–408. https://doi.org/10.1006/meth.2001.1262.

López-Tello, J., Pérez-García, V., Khaira, J., Kusinski, L.C., Cooper, W.N., Andreani, A., Grant, I., Fernández de Liger, E., Lam, B.Y., Hemberger, M., et al. (2019). Fetal and trophoblast PI3K p110α have distinct roles in regulating resource supply to the growing fetus in mice. Elife 8. https://doi.org/10.7554/eLife.45282.

Lu, Q., and Zhu, L. (2020). The Role of Semaphorins in Metabolic Disorders. Int J Mol Sci 21, E5641. https://doi.org/10.3390/ijms21165641.

Lu, Z.-Y., Cen, C., Shao, Z., Chen, X.-H., Xu, C.-F., and Li, Y.-M. (2016). Association between serum α-L-fucosidase and non-alcoholic fatty liver disease: Cross-sectional study. World J Gastroenterol 22, 1884–1890. https://doi.org/10.3748/wjg.v22.i5.1884.

Luche, H., Weber, O., Nageswara Rao, T., Blum, C., and Fehling, H.J. (2007). Faithful activation of an extra-bright red fluorescent protein in “knock-in” Cre-reporter mice ideally suited for lineage tracing studies. Eur J Immunol 37, 43–53. https://doi.org/10.1002/eji.200636745.

Montaigne, D., Butruille, L., and Staels, B. (2021). PPAR control of metabolism and cardiovascular functions. Nat Rev Cardiol 18, 809–823. https://doi.org/10.1038/s41569-021-00569-6.

Musial, B., Fernandez-Twinn, D.S., Vaughan, O.R., Ozanne, S.E., Voshol, P., Sferruzzi-Perri, A.N., and Fowden, A.L. (2016). Proximity to Delivery Alters Insulin Sensitivity and Glucose Metabolism in Pregnant Mice. Diabetes 65, 851–860. https://doi.org/10.2337/db15-1531.

Napso, T., Yong, H., Lopez-Tello, J., and Sferruzzi-Perri, A.N. (2018). The role of placental hormones in mediating maternal adaptations to support pregnancy and lactation. Front. Physiol. 9. https://doi.org/10.3389/fphys.2018.01091.

Napso, T., Zhao, X., Lligoña, M.I., Sandovici, I., Kay, R.G., George, A.L., Gribble, F.M., Reimann, F., Meek, C.L., Hamilton, R.S., et al. (2021). Placental secretome characterization identifies candidates for pregnancy complications. Commun Biol 4, 701. https://doi.org/10.1038/s42003-021-02214-x.

Ong, P.S., Wang, L.Z., Dai, X., Tseng, S.H., Loo, S.J., and Sethi, G. (2016). Judicious Toggling of mTOR Activity to Combat Insulin Resistance and Cancer: Current Evidence and Perspectives. Front Pharmacol 7, 395. https://doi.org/10.3389/fphar.2016.00395.

Petry, C.J., Evans, M.L., Wingate, D.L., Ong, K.K., Reik, W., Constância, M., and Dunger, D.B. (2010). Raised late pregnancy glucose concentrations in mice carrying pups with targeted disruption of H19delta13. Diabetes 59, 282–286. https://doi.org/10.2337/db09-0757.

Rieck, S., and Kaestner, K.H. (2010). Expansion of beta-cell mass in response to pregnancy. Trends Endocrinol Metab 21, 151–158. https://doi.org/10.1016/j.tem.2009.11.001.

Romero-Calvo, I., Ocón, B., Martínez-Moya, P., Suárez, M.D., Zarzuelo, A., Martínez-Augustin, O., and de Medina, F.S. (2010). Reversible Ponceau staining as a loading control alternative to actin in Western blots. Anal Biochem 401, 318–320. https://doi.org/10.1016/j.ab.2010.02.036.

Salazar-Petres, E.R., and Sferruzzi-Perri, A.N. Pregnancy-induced changes in β-cell function: what are the key players? The Journal of Physiology n/a. https://doi.org/10.1113/JP281082.

Sferruzzi-Perri, A.N. (2018). Assessment of Placental Transport Function in Studies of Disease Programming. Methods Mol. Biol. 1735, 239–250. https://doi.org/10.1007/978-1-4939-7614-0_14.

Simmons, D.G., Fortier, A.L., and Cross, J.C. (2007). Diverse subtypes and developmental origins of trophoblast giant cells in the mouse placenta. Dev. Biol. 304, 567–578. https://doi.org/10.1016/j.ydbio.2007.01.009.

Sparago, A., Cerrato, F., Vernucci, M., Ferrero, G.B., Silengo, M.C., and Riccio, A. (2004). Microdeletions in the human H19 DMR result in loss of IGF2 imprinting and Beckwith-Wiedemann syndrome. Nat Genet 36, 958–960. https://doi.org/10.1038/ng1410.

Stern, C., Schwarz, S., Moser, G., Cvitic, S., Jantscher-Krenn, E., Gauster, M., and Hiden, U. (2021). Placental Endocrine Activity: Adaptation and Disruption of Maternal Glucose Metabolism in Pregnancy and the Influence of Fetal Sex. Int J Mol Sci 22, 12722. https://doi.org/10.3390/ijms222312722.

Wood, A.J., and Oakey, R.J. (2006). Genomic imprinting in mammals: emerging themes and established theories. PLoS Genet 2, e147. https://doi.org/10.1371/journal.pgen.0020147.

Wu, C.-L., Zhao, S.-P., and Yu, B.-L. (2015). Intracellular role of exchangeable apolipoproteins in energy homeostasis, obesity and non-alcoholic fatty liver disease. Biol Rev Camb Philos Soc 90, 367–376. https://doi.org/10.1111/brv.12116.

Yamazawa, K., Kagami, M., Nagai, T., Kondoh, T., Onigata, K., Maeyama, K., Hasegawa, T., Hasegawa, Y., Yamazaki, T., Mizuno, S., et al. (2008). Molecular and clinical findings and their correlations in Silver-Russell syndrome: implications for a positive role of IGF2 in growth determination and differential imprinting regulation of the IGF2-H19 domain in bodies and placentas. J Mol Med (Berl) 86, 1171–1181. https://doi.org/10.1007/s00109-008-0377-4.

